# 16S rRNA sequence embeddings: Meaningful numeric feature representations of nucleotide sequences that are convenient for downstream analyses

**DOI:** 10.1101/314260

**Authors:** Stephen Woloszynek, Zhengqiao Zhao, Jian Chen, Gail L. Rosen

## Abstract

Advances in high-throughput sequencing have increased the availability of microbiome sequencing data that can be exploited to characterize microbiome community structure *in situ.* We explore using word and sentence embedding approaches for nucleotide sequences since they may be a suitable numerical representation for downstream machine learning applications (especially deep learning). This work involves first encoding (“embedding”) each sequence into a dense, low-dimensional, numeric vector space. Here, we use Skip-Gram word2vec to embed *k*-mers, obtained from 16S rRNA amplicon surveys, and then leverage an existing sentence embedding technique to embed all sequences belonging to specific body sites or samples. We demonstrate that these representations are biologically meaningful, and hence the embedding space can be exploited as a form of feature extraction for exploratory analysis. We show that sequence embeddings preserve relevant information about the sequencing data such as *k*-mer context, sequence taxonomy, and sample class. Specifically, the sequence embedding space resolved differences among phyla, as well as differences among genera within the same family. Distances between sequence embeddings had similar qualities to distances between alignment identities, and embedding multiple sequences can be thought of as generating a consensus sequence. Using sample embeddings for body site classification resulted in negligible performance loss compared to using OTU abundance data. Lastly, the *k*-mer embedding space captured distinct *k*-mer profiles that mapped to specific regions of the 16S rRNA gene and corresponded with particular body sites. Together, our results show that embedding sequences results in meaningful representations that can be used for exploratory analyses or for downstream machine learning applications that require numeric data. Moreover, because the embeddings are trained in an unsupervised manner, unlabeled data can be embedded and used to bolster supervised machine learning tasks.

**Author summary:** Improvements in the way genomes are sequenced have led to an abundance of microbiome data. With the right approaches, researchers use this data to thoroughly characterize how microbes interact with each other and their host, but sequencing data is of a form (sequences of letters) not ideal for many data analysis approaches. We therefore present an approach to transform sequencing data into arrays of numbers that can capture interesting qualities of the data at the sub-sequence, full-sequence, and sample levels. This allows us to measure the importance of certain microbial sequences with respect to the type of microbe and the condition of the host. Also, representing sequences in this way improves our ability to use other complicated modeling approaches. Using microbiome data from human samples, we show that our numeric representations captured differences between different types of microbes, as well as differences in the body site location from which the samples were collected.

## Introduction

Recent advances in high-throughput sequencing techniques have dramatically increased the availability of microbiome sequencing data, allowing investigators to identify genomic differences among microbes, as well as characterize microbiome community structure *in situ*. A microbiome is defined as the collection of microorganisms (bacterial, archaeal, eukaryotic, and viral) that inhabit an environment. Recent work has shown that there exists reciprocal interplay between microbiome and environment, such that the configuration of microbes is often influenced by shifts in environmental state (such as disease in human hosts [1–3], chemical alterations in soil [4, 5], or oceanic temperature changes [6]). Conversely, the environmental state may be impacted by particular microbial profiles [7–10].

Identifying and characterizing important microbial profiles often entails sequencing collections of heterogeneous, fragmented genomic material, which act as a proxy for the microbiome’s configuration in *situ*. Sequencing these fragments yields short strings of nucleotides (“sequence reads”) with no easily discernible clues to determine from which microbe they originated. Still, the order of nucleotides within each sequence provides enough information for approaches that utilize sequence alignment or sequence denoising algorithms, along with taxonomic reference databases, to quantify the abundance of sequences belonging to different microbes at different taxonomic levels (e.g., genus), which in turn can be associated with environmental factors [11, 12]. Thus, a given sample is represented as a vector of hundreds, often thousands, taxonomic counts (nonnegative integers).

An alternative approach to aligning or denoising nucleotide sequences is to represent the nucleotide sequences numerically. One can then search for similarities among these numeric features. In addition, these numeric representations are more suitable for machine learning algorithms. Two examples of such approaches are one-hot-encoding and *k*-mer (n-gram) counting [13, 14]. With one-hot-encoding (also referred to as generating binary indicator sequences [15]), each sequence is binarized - that is, each nucleotide (ACGT) is represented as a unit vector of length four, with a value of one indicating the presence of a particular nucleotide. *k*-mer counting, on the other hand, counts the frequency of all possible substrings of length *k* in a sequence. A larger *k* yields a higher dimensional, more sparse representation of the sequence since there are more possible *k*-mers (4^k^) (“the curse of dimensionality” [13, 16]), but a smaller *k* is unlikely to capture much of the nucleotide-to-nucleotide sequential variation among sequences [17]. Either set of engineered features (one-hot-encodings or *k*-mer frequencies) can then be used in various machine learning algorithms to characterize the sequences in some way [18–21]. Still, one-hot-encoding or *k*-mer frequencies, when *k* is large, yield sparse, high-dimensional features that often present difficulties during training [22]. In addition, neither approach encodes the relative ordering of the *k*-mers [13, 23].

A more suitable representation of nucleotide sequences involves first encoding (“embedding”) each sequence into a dense, numeric vector space via the use of word embedding algorithms such as word2vec [18]. Word embeddings are commonly used for natural language processing [18, 24–27]. Various architectures exist, but their objective is generally the same: capture semantic and lexical information of each word based on that word’s context – *i.e.,* its neighboring set of words. Each word is represented in a vector space of predefined length, where semantically similar words are placed near one another. Thus, *k*-mer representations of sequences could be embedded in such a way that their context is preserved (the position of *k*-mers relative to their neighbors), and they become suitable for down-stream machine learning approaches. Recent work has successfully embedded short, variable-length DNA *k*-mers [13], as well as protein sequences for down-stream tasks such as protein structure prediction [28]. Ng [13] showed that the cosine similarity between embedded *k*-mers is positively correlated with Needleman-Wunsch scores obtained via global sequence alignment. In addition, he showed that vector arithmetic of two *k*-mer embeddings is analogous to concatenating their nucleotide sequences. This finding is consistent with work demonstrating the ability of vector arithmetic to solve word analogies, such as “King is to Queen, as Man is to ______” [29].

Thus, here we explore word embeddings as a means to represent 16S rRNA amplicon sequences, obtained from microbiome samples, as dense, low-dimensional features that preserve *k*-mer context (*i.e.,* leverage the relative position of *k*-mers to their neighbors). We use Skip-Gram word2vec to perform the initial *k*-mer embedding. Then, we leverage an existing sentence embedding technique [30] to embed individual nucleotide sequences or sets of sequences (*e.g.,* all sequences belonging to a given sample) from *k*-mer embeddings. This sentence-embedding technique interestingly does not explicitly encode word order; yet, it has shown to outperform competing methods such as recurrent neural networks in textual similarity tasks [30].

The sentence embedding procedure is simple, but effective, consisting of down-weighting the embeddings for high frequency *k*-mers, averaging the *k*-mer embeddings that constitute a given sequence or set of sequences (forming a sequence or set embedding), and then subtracting the projection of the sequence/set embedding to its first principal component (“common component removal”, which, per [30], we refer to as “denoising”). Representing nucleotide sequences in vector space provides multiple benefits that may prove valuable in characterizing a microbiome: (1) the embeddings are dense, continuous, and relatively low-dimensional (compared to using *k*-mer frequencies, for example), making them suitable for various down-stream machine learning tasks; (2) they leverage *k*-mer context, yielding potentially superior feature representations compared to *k*-mer frequencies; (3) once trained, the *k*-mer-to-embedding mapping vectors can be stored and used to embed any set of 16S rRNA amplicon sequences; (4) the embedding model can be trained with data that are independent of the query sequences of interest, such that the training procedure can leverage a significant amount of unlabeled data that would otherwise go unused; (5) feature extraction is performed at the sequence level, enabling one to detect relationships between sample-level information (*e.g.,* soil quality) and all sequences belonging to a given sample and then traceback to determine not only which sequences are key, but also which *k*-mers; and (6) once important *k*-mers are identified, because the embedding initially takes place at the *k*-mer level, the *k*-mer contextual information is available, which may indicate the neighborhood noteworthy *k*-mers occupy.

In this work, we prove that the embedding space performs well at classifying samples, predicting the correct sample class (*e.g.,* body site) given the embedding of all its sequences. Moreover, because these embeddings are encoded from *k*-mer embeddings, their classification performance helps justify the use of *k*-mer embeddings as input in more complex architectures such as deep neural networks. We show that the embedding space provides a set of meaningful features that capture sample-level (sample class), taxonomic-level (sequence, read), and sequence-level (*k*-mer, *k*-mer context) characteristics, which not only justifies the use of the embedding space for supervised tasks such as classification, but also justifies its use for unsupervised feature extraction, to capture meaningful signal for the exploratory phase of a given analysis. Lastly, we illustrate approaches that may help disentangle what the embedding learned from the data, in the context of microbiome information, such as taxonomy and sample information.

## Results and Discussion

We will use the following terminology for the remaining sections. “Query” sequences and *k*-mers are all sequences and *k*-mers that were not used for training (and hence are not in the GreenGenes reference database). A “*k*-mer” embedding is an d-dimensional vector containing the embedding of a nucleotide subsequence of length *k*. A “sequence embedding” is a d-dimensional vector formed by performing the sentence embedding approach by [30] on all *k*-mers belonging to a single nucleotide sequence (*e.g.,* a 16S rRNA full-length sequence or read). “Cluster embeddings” or “sample embeddings” are also d-dimensional vectors, but they encode an embedding that includes all *k*-mers belonging to a set of sequences (such as a set identified via clustering) or a set that includes all sequences belonging to a single sample, respectively. Lastly, “denoising” refers to common-component removal (not dada2-style “sequence denoising algorithms”), which is performed when we apply the sentence embedding approach to sequences, sequence reads, clusters of sequences, or all sequences belonging to specific samples.

### Evaluation of sequence embeddings on full-length 16S rRNA amplicon sequences

We began by evaluating the performance of sequence embeddings. DNA alignment combined with clustering [11] or sequencing denoising algorithms [12] are readily capable of identifying sequence-level differences between genera. We consequently aimed to discern if genus-level differences were in fact detectable in the sequence embedding vector space. Genus level resolution is not only relevant to characterizing a microbial community; it also would suggest that the vector space is capable of capturing subtle, but important sequence differences among taxa. These differences may be critical in characterizing data at the sample level, particularly during classification, where its desired to discern how the configuration of microbes comprising a sample (*i.e.,* all its sequences) is influenced by sample-level information such as soil quality.

The *k*-mer embedding space was obtained by training Skip-Gram word2vec on 2,262,986 full-length 16S rRNA amplicon sequences from the GreenGenes reference database (see S1 Appendix for parameter selection). An independent set of 16,699 16S rRNA sequences from the KEGG REST server [31] were obtained as our test dataset. (We will refer to these sequences as “KEGG 16S sequences” henceforth.) 14,520 contained *k*-mers that intersected with the training set and thus were sequence-embedded using the sentence embedding approach by [30]. Briefly, for each sequence of length N, its *N* – *k* +1 *k*-mers were embedded into vectors of length d. With these *k*-mer embeddings, we calculated a weighted average by summing across all *k*-mer embeddings (element-wise), down-weighting high-frequency *k*-mers, and dividing by the total number of *k*-mers (*N* – *k* + 1). The sequence embedding was then obtained b subtracting its projection to its first principal component (denoising).

### Taxa separate in the sequence embedding space at phylum and genus levels

To visualize the sequence embedding space, we performed dimensionality reduction with t-distributed stochastic neighbor embedding (t-SNE) [32]. Fig 1 shows the 2-dimensional projections of 256-dimensional sequence embeddings that were formed by averaging 10-mer (*k*-mer where k=10) embeddings. Shown are the eight most abundant phyla (A). Sequences (points) grouped as a function of their phylum classification. When we focus on genus classifications within a family, grouping persists. For example, within Bacillaceae (B, top), sequences from *Geobacillus spp*. (blue) separated from *Bacillus spp*. (red); within Enterobacteriaceae (B, middle), sequences from *Yersinia spp*. (red) separated from *Klebsiella spp*.; and within Streptococcaceae (B, bottom), sequences from *Streptococcus spp*. separated from *Lactococcus spp*. These observations suggest that the sequence embedding preserves genus-level resolution.

**Fig 1.**
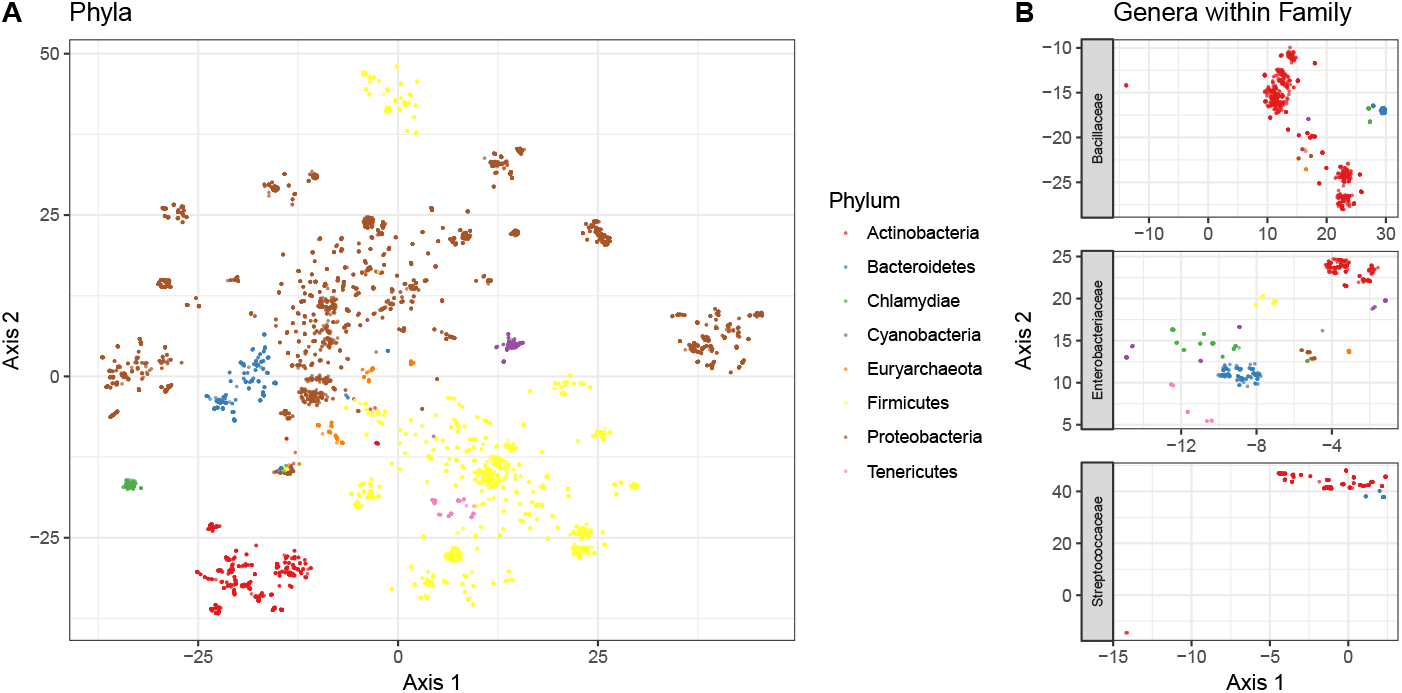
t-SNE projection of sequence embeddings from KEGG 16S sequences. Embedding results were generated using 256 dimensional embeddings of 10-mers that were denoised. A: 2-dimensional projection via t-SNE of the sequence embedding space from 14,520 KEGG 16S sequences. The position of each sequence (points) are colored based on their phylum designation. B: t-SNE projection of sequences that belong to different genera within the same family.

### Pairwise cosine similarity between sequence embeddings agrees with pairwise sequence alignment identity

With the 2,962 sequence embeddings shown as points in Fig 1B, we obtained their nucleotide sequences and performed pairwise global sequence alignment via v-search [33]. These sequences belonged to Bacillaceae, Streptococcaceae, and Enterobacteriaceae families. Our objectives were to (1) determine, for a pair of sequences, if the distance (cosine similarity) between their sequence embeddings was equivalent to the distance (alignment score) between their nucleotide sequences and (2) quantify the variation in which distances between sequence embeddings vary as a function of taxon. We expected that because sequences are more similar among closely related taxa (such as taxa found within the same genus), the average pairwise cosine similarity between sequences in a lower taxonomic level should be larger. Fig 2 shows the (z-scored) within-taxon distributions of (1) pairwise cosine similarity between sequence embeddings and (2) pairwise percentage of identity between nucleotide sequences using v-search. The results suggest that sequence embeddings behave similarly to sequence alignment in terms of pairwise distance, and as we move down the taxonomic hierarchy, the distance z-scores increased by 0.629 (*p* < 0.001, *R*^2^ = 0.795) and 0.636 (*p* < 0.001, *R*^2^ = 0.803) standard deviations for sequence embeddings and v-search, respectively. Moreover, within genus, the standard deviation of scores for either approach (genus sequence embedding sd=0.415; genus v-search alignment sd=0.399) is similar, suggesting the spread of distances between sequences from the same genus is equivalent between approaches. Therefore, this evidence shows that sequence embeddings are similar to global alignment at distinguishing intra-genus differences among sequences.

**Fig 2.**
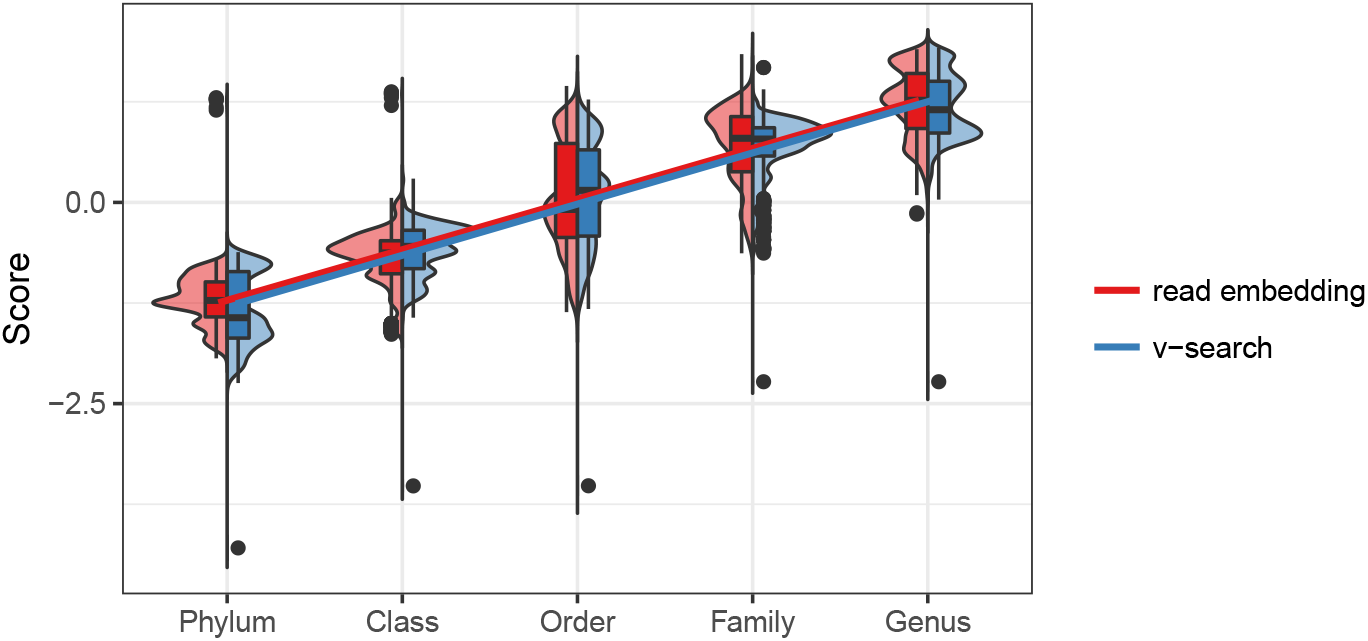
Within-taxon distribution of pairwise sequence alignment similarity verses pairwise embedding similarity. Embedding results were generated using 256 dimensional embeddings of 10-mers that were denoised. For a given taxonomic level, the red violin plots represent the distribution of pairwise cosine similarity between all sequence embeddings from the 14,520 KEGG 16S rRNA sequences, whereas the blue violin plots represent the distribution of pairwise nucleotide sequence identity using global alignment via v-search. Both sets of scores were z-scored to make them visually comparable. Linear regression best fit lines are shown to ease interpretation.

### Embedding clusters of sequences is comparable to calculating their consensus sequence

Given that the sequence embeddings behaved analogously with alignment, we hypothesized that a sequence embedding was equivalent to performing global sequence alignment and calculating a consensus *k*-mer sequence (the majority nucleotide in each column of a global alignment). By extension, averaging together multiple sequence embeddings could be thought of as calculating a consensus sequence of their nucleotide sequences. To verify this, we used v-search to cluster the 14,520 KEGG 16S sequences and then obtained the consensus sequence from each of the resulting 176 clusters. We calculated (1) the sequence embedding for each consensus sequence and (2) the cluster embedding for all sequences within a cluster (*i.e.,* a weighted mean sequence embedding for a given cluster, but the projection to the first principal component removed only after the complete cluster embedding is constructed). We expected that if an embedding of multiple sequences was in fact similar to calculating the consensus sequence of their nucleotide sequences, then the cosine similarity would be largest between the sequence embedding of a cluster’s consensus sequence and its cluster embedding. The cosine similarities between the consensus sequence embeddings and each cluster embeddings are shown in Fig 3. The largest cosine similarity occurs on the diagonal, between consensus and cluster embeddings pertaining to the same cluster. This observation suggests that an embedding of multiple sequences can be viewed as a consensus sequence.

**Fig 3.**
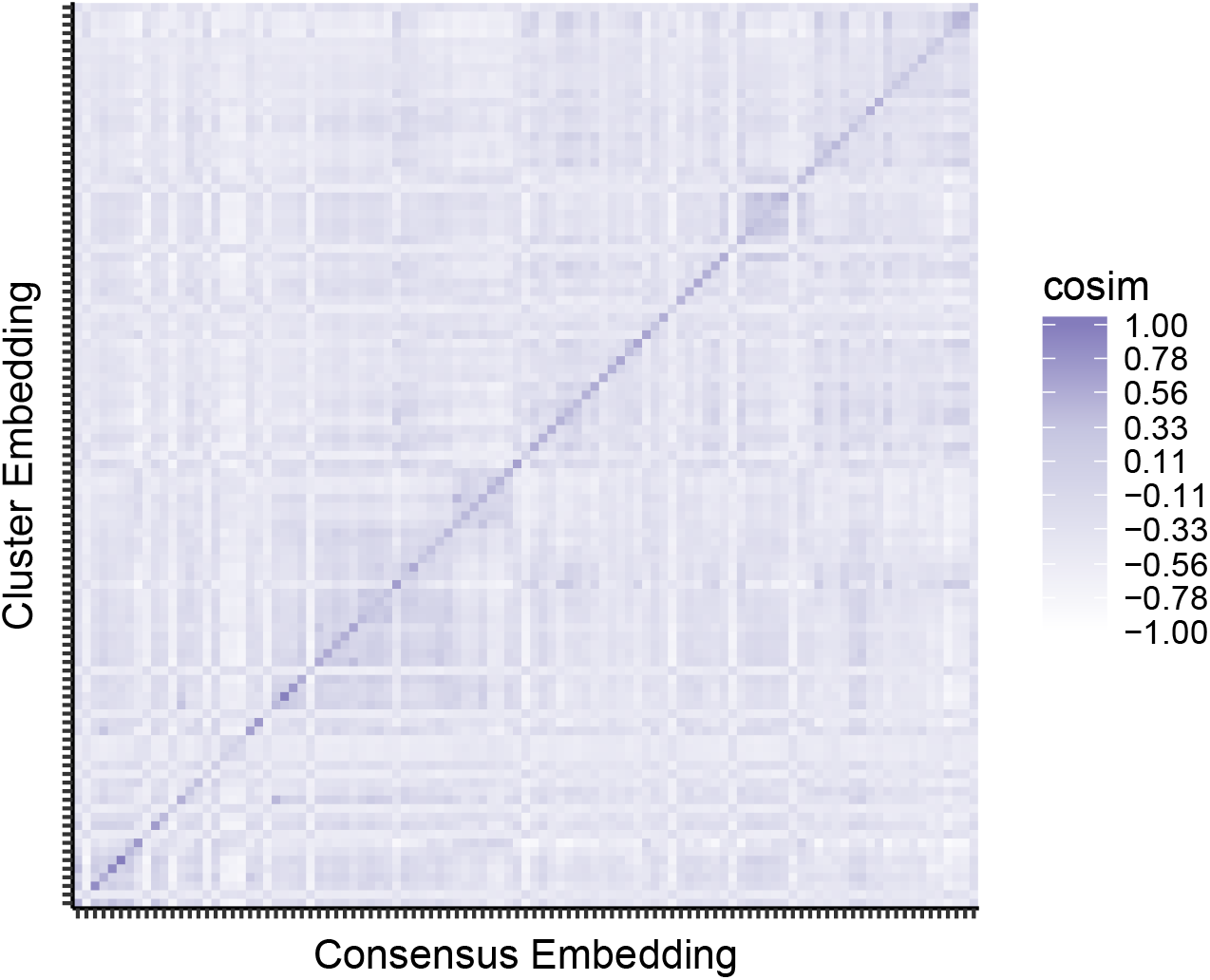
Agreement between consensus sequence embeddings and their cluster embeddings. For each cluster, all KEGG 16S sequences were embedded into a cluster embedding, and the cluster’s v-search consensus sequence was embedded into a consensus embedding. The pairwise cosine similarities between all consensus and cluster embeddings are shown. They are sorted based on the (arbitrary) index for cluster membership. Darker shading indicates larger cosine similarity betweens cluster and consensus embeddings. Thus, the dark diagonal represents that the cluster embedding is similar to the consensus embedding for that cluster.

### Clustering the embeddings yield higher fidelity clusters for species than for higher taxonomic levels

To better understand how well the embedding space represented sequence data, particularly in its ability to resolve subtle differences among sequences in terms of some similarity statistics, we clustered the KEGG 16S sequence embeddings. We used *K*-means [34], which iteratively clusters sequence embeddings into the closest centroid and updates the centroid based on each cluster it contains. The free parameter K was chosen according to the total number of taxa in a given taxonomic level. For example, at the species level, there were 385 unique taxa; thus, we set K to 385, which resulted in 385 *K*-means generated clusters.

Table 1 shows the clustering performance for one of the embedding models (10-mers, 256 dimensions, denoised). *K*-means performed best at the species level for all metrics. For higher taxonomic levels, performance declined. This shows that the sequence embedding preserved short- and medium-term distance better than long-term distance — that is, similar sequences remain close in the embedding space. While the similarity between sequences belonging to different phyla is not well preserved in the embedding space, the clustering performance still suggests that the sequence embeddings are informative and capable of preserving the sequence similarity among nucleotide sequences.

**Table 1.** Clustering analysis of KEGG seauence embeddings.

| Taxon | K | P | C | O | F | G | S |
| --- | --- | --- | --- | --- | --- | --- | --- |
| # of Taxa | 2 | 35 | 77 | 143 | 281 | 697 | 385 |
| K | 2 | 35 | 77 | 143 | 281 | 697 | 385 |
| Homogeneity | 0.11 | 0.88 | 0.94 | 0.94 | 0.96 | 0.97 | 0.98 |
| Completeness | 0.02 | 0.42 | 0.56 | 0.78 | 0.79 | 0.83 | 0.94 |
| ARI | -0.01 | 0.24 | 0.32 | 0.36 | 0.43 | 0.47 | 0.84 |
| AMI | 0.02 | 0.42 | 0.55 | 0.66 | 0.75 | 0.76 | 0.91 |
| NMI | 0.04 | 0.61 | 0.72 | 0.80 | 0.87 | 0.90 | 0.96 |

### Evaluation of sequence embeddings on 16S rRNA amplicon sequences from the American Gut Project

Next we aimed to try our embedding approach on empirical data. We obtained sequencing data of microbiota from three body sites sequenced by the American Gut project [35]. After preprocessing, 11,341 samples from each of three body sites (fecal, skin, oral) were embedded. Unlike the KEGG 16S sequences described above, the fact that these are reads (and not full-length 16S rRNA sequences) presents a new challenge in that they are significantly shorter, spanning only 125 nucleotides per read; thus, each read is composed of, at most, 116 *k*-mers for a 10-mer embedding. The *k*-mer, sequence, and sample embedding spaces are shown in Fig 4. There was clear grouping among samples from body sites (Fig 4C) and phyla for sample and sequence embeddings, respectively (Fig 4B). For the *k*-mer embedding, discerning any meaningful patterns is more difficult since the embedding simply encodes contextual information for each *k*-mer. We were interested in the relative position of single nucleotide differences for a given k-mer. In Fig 4A, we mark the position of every 10-mer present that differs from AAAAAAAAAA by one nucleotide. As expected, despite subtle differences among the 10-mers, their relative position in the embedding space is broad, suggesting that each 10-mer’s context significantly influenced its embedding.

**Fig 4.**
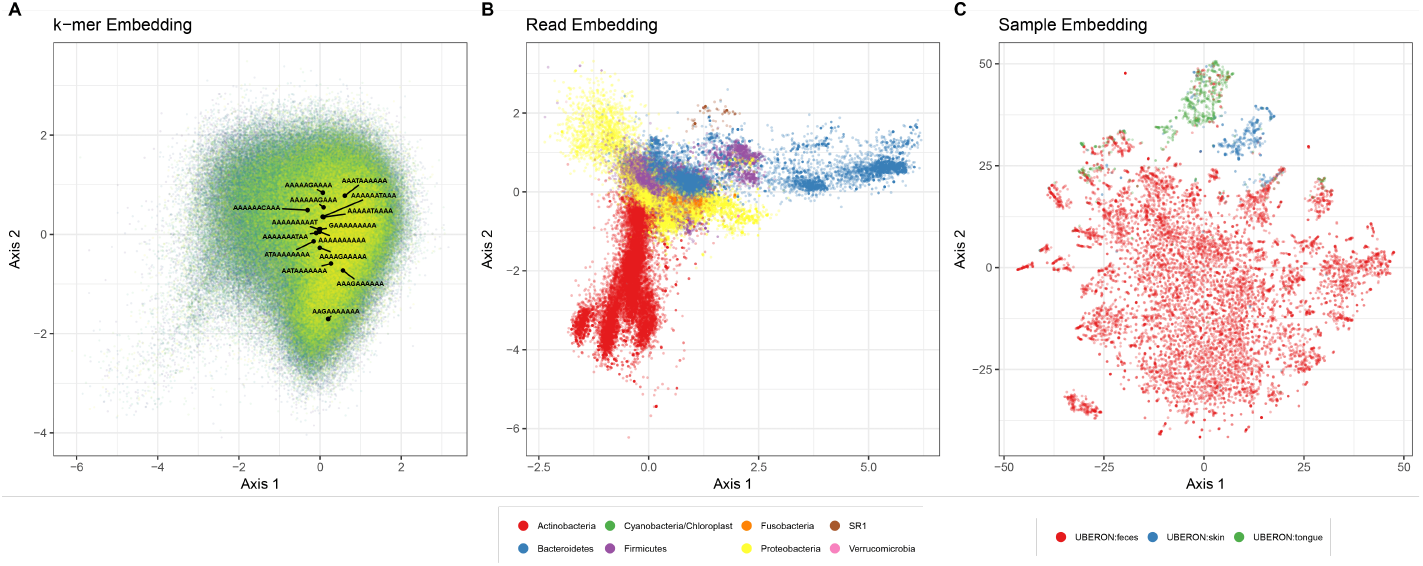
Lower dimensional projections of k-mer, sequence, and sample embeddings. Embedding results were generated using 256 dimensional embeddings of 10-mers that were denoised. A: A 2-dimensional projection via independent component analysis of the 10-mer embedding space from the GreenGenes training sequences. 406,922 unique 10-mers are shown. The position of 10-mers that differ by one nucleotide from AAAAAAAAAA are labeled to demonstrate that it is not simply sequence similarity that is preserved, since these sequences span a wide range in the embedding space. The *k*-mers were sorted alphabetically and ranked; the alphabetical progression of the indexes are shaded from yellow to green. B: A 2-dimensional projection via independent component analysis. 705,598 total sequences embeddings from 21 randomly chosen American Gut samples (7 from each class) are shown. The position of each sequence (points) is colored based on its phylum designation (only the 7 most abundant phyla are shown). C: 2-dimensional t-SNE projection of the 11,341 American Gut sample embeddings. The position of each sample (points) is colored based on its body site label.

### Sample embeddings show negligible performance loss compared to OTU abundances for classification

We evaluated testing performance using 256-dimensional sample embeddings as features to classify body site (fecal, skin, oral). We used a multinomial lasso classifier [36, 37] to obtain a sparse set of regression coefficients that we could use to interrogate predictive nodes at both sequence and *k*-mer embedding levels. We compared generalizability (testing performance) of five different sets of features: (1) QIIME-generated [11] OTU abundances (centered-log-ratio (clr) transformed [38, 39]), (2) the top-256 principal components after applying principal component analysis (PCA) to the OTU abundances, (3) 1000 pseudo-OTUs (clr transformed) that were generated by clustering sequence embeddings, (4) 256-dimensional sample embeddings generated with 6-mers, and (5) 256-dimensional sample embeddings generated with 10-mers (Table 2). The PCA approach provided us with an alternative form of dimensionality reduction (to ensure the dimensionality reduction of the embedding does not yield an unfair advantage compared to the OTU abundances), whereas the rationale behind generating pseudo-OTUs was to more explicitly represent sequence abundances compared to averaging sequence embedding vectors, which yields a consensus sequence embedding that is weighted as a function of sequence frequency. Besides the PCA approach, all models performed well, with OTU abundances performing best (0.977 mean balanced accuracy). The PCA approach failed to predict “skin” for any sample and rarely predicted “tongue,” leading to 0.317 mean balanced accuracy. Pseudo-OTUs and 10-mer sample embeddings performed comparably, with mean balanced accuracies of 0.968 and 0.962, respectively. This suggests that generating sample embeddings yields predictive features that remain generalizable despite drastically reducing the dimensionality of the data. It is also clearly a superior dimensionality reduction approach compared to applying PCA to an OTU abundance table.

**Table 2.**
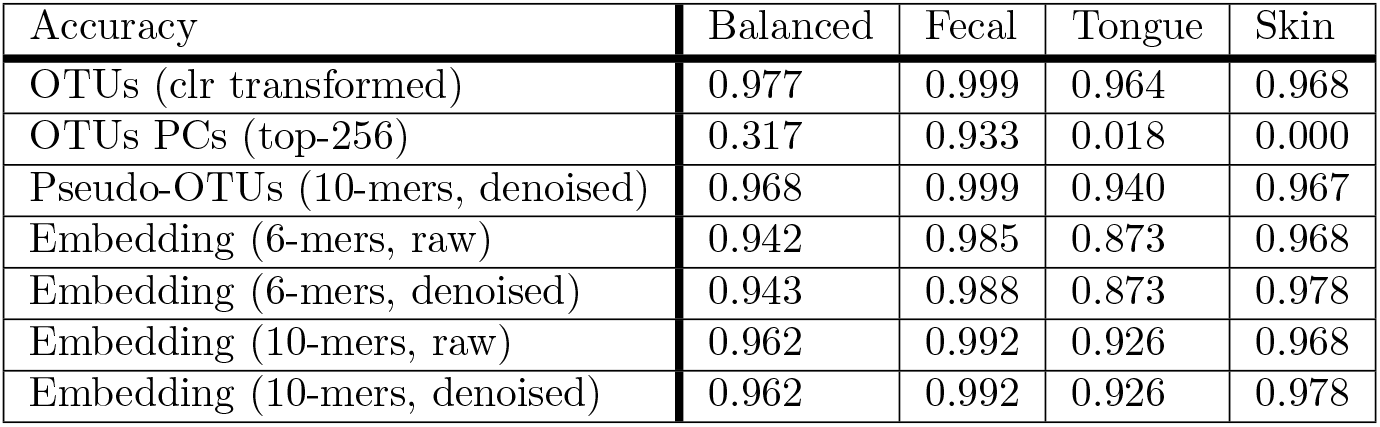
Sample embedding classification performance.

### Specific sequences activate specific nodes and these activations are a function of the sequence’s taxon and the sample’s body site

We next aimed to determine (1) how the sample embedding matured as sequence embeddings from different taxa were introduced and (2) how specific dimensions (nodes) in the embedding space (the hidden layer) influenced the final sample embedding. When simply optimizing for classification performance, it is not unreasonable to treat a neural network, or any machine learning model, as a black box, but with a better understanding of what these models are learning, the potential for novel applications increases. Hence, the interpretation of the hidden layers in neural networks is an active area of research [26, 40–43]. One approach is to train a model and then traceback from its prediction to identify key features influential in the classification decision [40, 44]. By performing classification with the sample embedding treated as features, we can traceback to disentangle how key nodes represented the embedding (*e.g.,* whether particular nodes activated as a function of taxon), as well as discern the impact specific reads, and hence taxa, had on both the sample embedding and classification.

We obtained the same sparse set of regression coefficients that were estimated via lasso to classify body site using sample embeddings of the American Gut data (see above). We calculated read activations (the linear combination of the lasso regression coefficients and the sequence embeddings) and then inspected how the cumulative sum of these activations affected the final body site classification (fecal, tongue, or skin).

Fig 5 shows the trajectory of the classification decision for a single tongue sample that was misclassified as fecal. The sequences were sorted and added to the embedding in order, based on their taxonomic classification (independently obtained via the naive Bayes RDP classifier). Fig 5A-B) show the trajectory of all nodes (their sum) in the embedding, and thus represents the overall classification decision as reads are introduced. Fig 5C-F show the activations for individual nodes that greatly impacted the classification decision (*i.e.,* they had large activations). Note that only Fig 5A shows the cumulative trajectory; Fig 5B-F show the activations specific for a given read (index).

**Fig 5.**
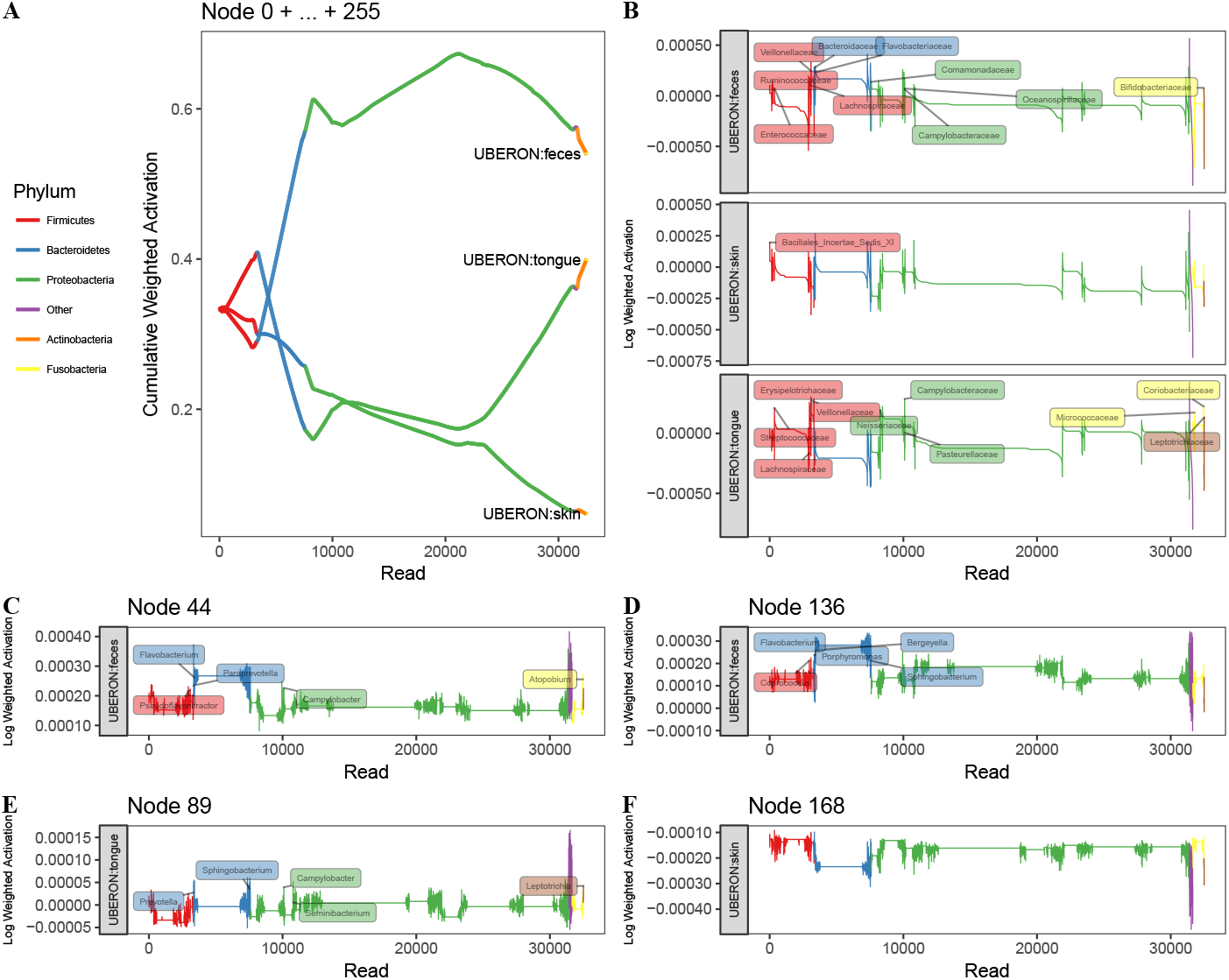
Maturation of the body site classification decision as sequence embeddings are introduced. Embedding results were generated using 256 dimensional embeddings of 10-mers without denoising (since the number of reads varies throughout the figure). One American Gut tongue sample is shown, which was misclassified by lasso as “fecal.” Read activations are defined as the linear combination of a given sequence embedding and the regression coefficients obtained from lasso for a particular body site. A: The trajectory of the body site classification decision via multinomial lasso. The cumulative activation is the sum of all read activations across all nodes (dimensions of the embedding) up until the introduction of a specific read. A body site is favored when it has the largest cumulative activation of the three body sites. Reads were sorted and introduced based on their taxon (phylum designations are color coded). B: The (non-cumulative) activations (across all nodes) for body site as reads are introduced (with no accumulation from previous reads). Genera labels are shown for reads with large activations. C-F: The (non-cumulative) activations for individual nodes and specific body sites as reads are introduced.

The initial set of reads introduced to the sample embedding were 3327 reads belonging to the phylum Firmicutes. This set of reads drove the classifier to favor a “skin” classification. Highly influential genera among these reads include *Granulicatella, Gemella,* and *Streptococcus,* which is consistent with work linking these genera to microbiota inhabiting the nares [45], as well as work demonstrating shifts in *Gemella* and *Streptococcus* occurring in response to atopic dermatitis [46]. As 4263 Bacteroidetes reads were introduced to the sample embedding, the classifier began to favor a “feces” classification. However, influential reads predominantly belonged to Flavobacteriaceae, a family often associated with oral microbiota [47, 48]. Perhaps during training, the encoding of this family became associated with k-mers belonging to gut sequences. Thus, the fact that the classifier incorrectly associated these oral microbiota with the fecal body site likely influenced the classifier to classify the sample incorrectly. The introduction of Proteobacteria influenced the classifier differently at different stages, depending on the type of Proteobacteria introduced. Initially, reads assigned to the family Comamonadaceae (x-axis read index=7591), a bacterial family associated with gut microbiota [49], reinforced the fecal classification. Favorability towards tongue increased upon introduction of reads (read index=8641) belonging to the genera *Neisseria,* and to a lesser extent, *Campylobacter* and *Haemophilus,* all of which have been linked to oral microbiota [47, 48]. *Acinetobacter* reads (read index=10866) that were favorable to a fecal classification were then introduced. *Acinetobacter* has been linked dental plaque accumulation, nosocomial respiratory disease, poor cholesterol profiles, gut dysbiosis, and colorectal cancer [50–53]; thus, *Acinetobacter*’s association with both oral and gut microbiota may have also contributed to the misclassification. Finally, *Pseudomonas* and *Stenotrophomonas* reads (read index=21189), as well as a short-lived contribution of *Atopobium, Rothia,* and *Actinomyces* reads from the phylum Actinobacteria (read index=31655) shifted the classifier closer to a tongue classification, which is consistent with literature [5, 48]. However, these oral shifts were unable to outweigh the misclassifications from the Flavobacteriaceae, and thus “fecal” was the final classification decision.

When we focused on individual nodes (dimensions in the embedding space), we were able to link nodes to specific taxa. For example, nodes 44 and 136 received non-zero regression (lasso) coefficients for classifying samples as fecal. These nodes had large activations for sequences stemming from *Prevotella* and *Flavobacterium* from the Bacteroidetes phylum, but node 136 was specific for large Bacteroides activations (sequence index=3328-3336). Node 168, on the other hand, was important in skin classifications (non-zero regression coefficients) and had relatively large activations for sequences from Firmicutes phyla, specifically *Staphylococcus*, *Gamella*, *Oribacterium*, *Veillonella*, and *Granulicatella*.

### *k*-mers associated with Lachnospiraceae genera were specific for different body sites and specific regions of the 16S rRNA gene

Because the embedding was trained with *k*-mers, and each sequence or sample embedding therefore amounts to a weighted sum of *k*-mer embedding vectors, we found it necessary to traceback further to the encoded *k*-mer information. This is similar to the question found in natural language processing where feature interpretation is necessary to elucidate how the neural network represents saliency and compositionality [26]. Trying to extract meaning from sequences of nucleotides is obviously a different question, but nevertheless, much can be learned from natural language processing in understanding the context of a given *k*-mer, its co-occurrence profile, and how sequential sets of *k*-mers influence a classification decision differently than their constituent parts. For example, recent work has attempted to identify regulatory regions in genomic sequences in this manner [41].

Thus, our objectives were to identify (1) *k*-mers that strongly influence the performance during sample-level (body site) classification and (2) characteristics of the sequences that contain these *k*-mers, such as the sequence’s taxonomic classification vis-á-vis the neighborhood in which important *k*-mers occur. We again obtained the sparse set of regression coefficients that were estimated via lasso to classify body site using sample embeddings of the American Gut data. We calculated *k*-mer activations (the linear combination of the lasso regression coefficients and the *k*-mer embeddings) and then selected the top-1000 *k*-mers with largest activations for classifying skin, tongue, or fecal body sites. These *k*-mers, when present in a sequence, have the single greatest influence on the classification outcome for a given sample, although it should be noted that the final classification decision is predominantly driven by high-frequency (after weighting) *k*-mers with relatively large activations.

Here we focus on the classification of skin samples. For each of the 3 sets (skin, tongue, fecal) of 1000 *k*-mers with large activations, we identified which reads from American Gut skin samples contained these *k*-mers. From the 11,838,849 reads in the 282 skin samples, 241,006 reads contained any of the top-1000 *k*-mers with large activations for skin, whereas 74,240 and 197,505 reads contained any of the top-1000 *k*-mers with large activations for tongue and fecal classification, respectively. Only 2781 reads contained *k*-mers from both skin and fecal-associated *k*-mers (the intersection), whereas 5056 reads contained both skin and tongue-associated *k*-mers. Also, 626 of the 2781 reads with both skin and fecal-associated *k*-mers belonged to only two samples (221, 405). Interestingly, the sample with 221 reads that contained skin- and fecal-associated *k*-mers was misclassified by the lasso classifier (using the parameterization optimized during training), but any general trend between misclassification and the proportion of high-activation *k*-mers was not apparent. This finding is consistent with how the embedding activations affect classification, as we demonstrated above; *k*-mer activations incrementally influence the classifier, with no single *k*-mer embedding having substantial impact on the ultimate classification decision. Still, as we have shown, the *k*-mer profiles associated with particular body sites are distinct.

To further characterize the *k*-mer profiles, we associated high-activation *k*-mers with the neighborhood of their read from which they originated. For each set of reads that contained the high-activation *k*-mers, we randomly sampled 25,000 reads and performed multiple alignment. Fig 6 shows the mapping regions in the multiple alignment for *k*-mers with large activations. We only show genera belonging to the family Lachnospiraceae because focusing on genera that share a family, for example, may present interesting patterns as to how specific *k*-mers, associated with specific taxa, favor specific regions in the 16S rRNA gene. The cluster of 6 *k*-mers (index 1-6) (TTTCGGAACT, CCAGAACTGA, TTTGGAGCTA, ACTTATAAAC, AACTGTTGCT, GAAACCGTGC) that span nucleotide 500 in the multiple alignment mapped to reads from *Catonella* (120 reads), *Clostridium* (59), *Oribacterium* (27), *Stomatobaculum* (101), and two unclassified genera (340). For each of these 6 *k*-mers, we calculated their nearest *k*-mer neighbors — that is, all *k*-mer embeddings with cosine similarity greater than 0.25 (arbitrarily chosen). TTTGGAGCTA and GAAACCGTGC shared ten neighbors, whereas TTTCGGAACT, CCAGAACTGA, and ACTTATAAAC shared one, suggesting that some context (neighboring *k*-mers) was shared among these *k*-mers. This helps confirm the positional relationship seen in Fig 6. Coupling these results with our observation that intra-family genera separate in the embedding space (Fig 1), this region of the multiple alignment may pinpoint a key location of the 16S rRNA variable region that distinguishes these taxa. While we do acknowledge this claim garners additional evidence, we do feel it is worthy of future work, particularly with datasets that can yield more trustworthy taxonomic assignments. In addition, another interesting avenue for future work may involve using full length 16S rRNA sequences to identify associations between *k*-mer and sequence embeddings to elucidate which variable regions are favored by specific taxa.

**Fig 6.**
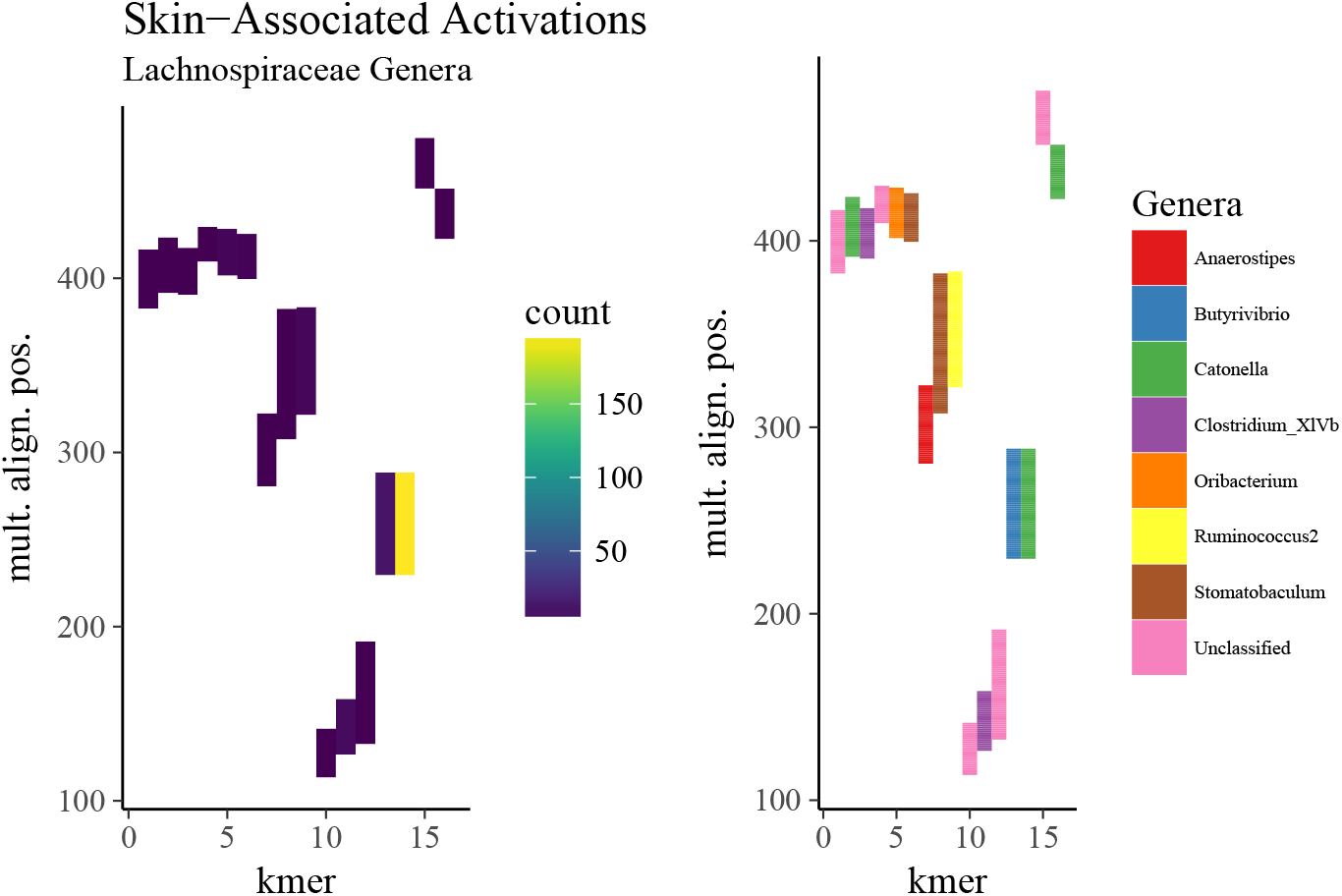
Regions within Lachnospiraceae reads among in which skin-associated *k*-mers mapped. The top-1000 *k*-mers with the largest activations (the linear combination of the *k*-mer embedding and regression coefficients obtained via lasso) for skin were identified. A random sample of 25,000 reads (from skin samples) containing these *k*-mers underwent multiple alignment. Shown is the relative position of *k*-mers found only in Lachnospiraceae reads from skin samples. The position of a given *k*-mer in the alignment spans its entire starting and ending position, including gaps. The frequency in which these *k*-mers mapped to positions (left column) in the multiple alignment were quantified (y-axis). The order of the *k*-mers in the heatmap (x-axis) was obtained via hierarchical clustering (Ward’s method) on Bray-Curtis distances. The Lachnospiraceae genus that most frequently occurred at a given alignment position for a given *k*-mer is colored (right column).

## The embedding corpora

### Many *k*-mers in the query datasets lacked embeddings

The total number of unique 6-mers and 10-mers in the GreenGenes reference database, used for training, was 4096 and 413,329, respectively. Of the 406,922 unique 10-mers in the KEGG 16S sequences, 270,676 were present in GreenGenes and thus had embeddings (*i.e.,* were present in the training set), whereas all *k*-mers were present for 6-mers. Of the 1,048,576 unique 10-mers in the American Gut reads, only 413,329 were embedded. Thus, the sample embeddings extracted meaningful features and maintained high performance despite over half of the American Gut *k*-mers not being used. This begs the question: what proportion of a given dataset is required to generate a meaningful embedding? Future work should explore (1) how sensitive the embedding space is to the degree of overlap between the of training and query vocabularies and (2) whether the requirement for sufficient overlap changes when the query data consists of longer sequences.

### Denoising sequence and sample embedding spaces removes mean background signal, which helps resolve the pairwise similarity among embeddings

The predominate effect of the denoising approach (removal of the projection to the first principal component) can be seen in Fig 7. Which shows the distribution of pairwise cosine similarity scores for *k*-mer, sequence, and sample embeddings using 6-mers and 10-mers. The results indicate three major trends. First, shorter *k*-mers result in larger pairwise cosine similarity between sequences, which is likely due to larger sequences being less common and, by extension, having more distinctive *k*-mer neighborhoods. Second, as more sequences are introduced to a given embedding, such as building a sample embedding compared to building a sequence embedding, the pairwise cosine similarity between sequences increases. We posit that as the number of sequences increases in an embedding, common *k*-mer embeddings begin to dominate in the same way that stop-words such as “the” may affect sentence embeddings. This, in turn, would explain the third trend: denoising mitigates this effect by removing the most common component that captures a mean background signal. Thus, we can conclude that denoising is pertinent in tasks involving cosine similarity where resolving subtle differences between sequences is paramount. Moreover, as the number of sequences introduced increases, denoising becomes more important.

**Fig 7.**
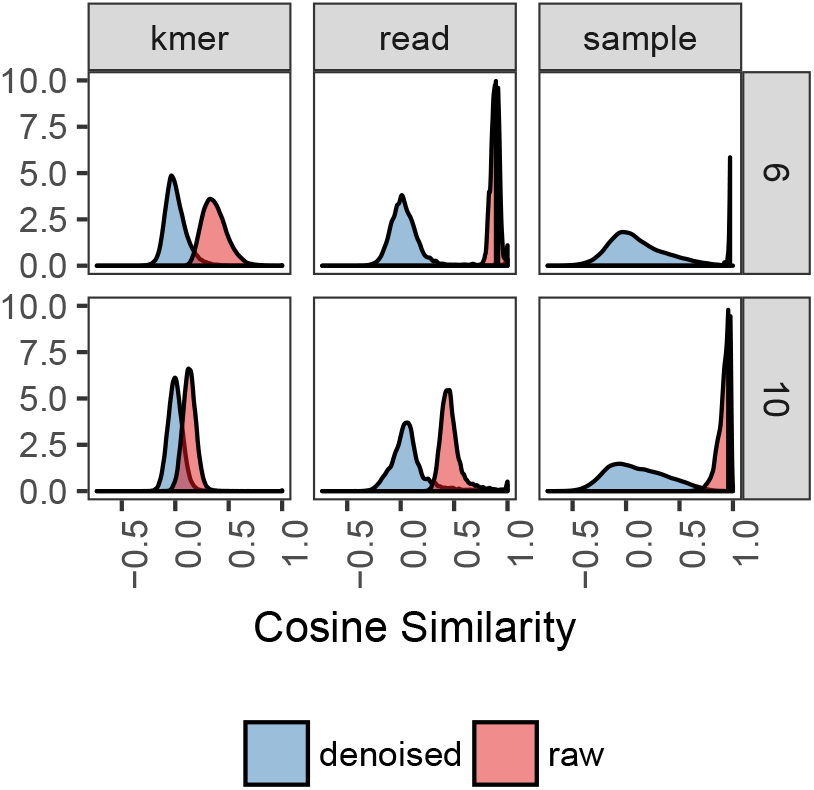
Distribution of pairwise cosine similarity for k-mer, sequence, and sample embeddings of the American Gut data. sequence and sample embeddings were calculated from 21 randomly selected (7 for each body site) samples. *k*-mer embeddings were those used during training with the GreenGenes sequences. Shown are distributions of pairwise cosine similarities as a function of k-mer size (rows), denoising (blue versus red), and embedding space (columns).

## Conclusion

Here we have applied word and sentence embedding approaches to generate *k*-mer, sequence, and sample embeddings of 16S rRNA amplicon sequencing data. We present our results at a time when deep neural network approaches are readily being applied to genomic sequencing data [54]. Although these approaches have been utilized to a lesser extent in microbiome research, increased use is likely inevitable as sequencing data becomes more available. Thus, obtaining meaningful numeric representations of microbiome sequences that does not suffer from the curse of dimensionality and can act as input to various machine learning architectures is necessary.

Our work demonstrates that sequence and sample embeddings are dense, lower-dimensional representations that preserve relevant information about the sequencing data such as *k*-mer context, sequence taxonomy, and sample class. We have shown that these representations are biologically meaningful and hence the embedding space can be exploited as a form of feature extraction for the exploratory phase of a given analysis. The sequence embedding space performs well compared to common approaches such as clustering and alignment, and the use of sample embeddings for classification seemingly results in little-to-no performance loss compared to the traditional approach of using OTU abundances. Because the sample embeddings are encoded from k-mer embeddings, its classification performance justifies further inquiry as pretrained input for more complex machine learning architectures such as deep neural networks. In addition, future work should aim at elucidating the effects of different training datasets and obtaining a better understanding of the feature representations at the *k*-mer level.

## Materials and Methods

### Primer on word2vec

Word2vec represents words as continuous vectors based on the frequency of pair co-occurrence in a context window of fixed length. We can understand it as mapping individual words (*k*-mers in our application) to points in a continuous, higher-dimensional space, such that words with similar semantic meaning are closer to one another.

Word2vec is a shallow, fully connected neural network with one hidden layer. It is unsupervised in the sense that labels need not be provided. There are two types: (1) Skip-Gram, where a word predicts its context (*i.e.,* neighboring words) and (2) continuous bag of words, where a context predicts its word. For Skip-Gram, the input and output layer have the same number of nodes as there are words in the vocabulary. For the input layer, given a vocabulary of length V, a single word w is encoded as a one-hot-vector of length V, where all elements in the vector are zero except the position corresponding to the target word w. The output layer is a vector of length V that contains the probability a randomly sampled word ¬*w* is w, which is obtained with a softmax function. The hidden layer is of size *V* × *d*, where d dictates the dimensionality of the embedding space and is much smaller in size than the vocabulary: *d* << *V*. Each row in the hidden layer corresponds to the embedding of some word w.

Training is accomplished via negative sampling to accelerate the training process. Instead of updating the hidden layer for the entire vocabulary each pass, for a given word w, a sampled subset of negative example words along with w are selected. The rows of the hidden layer corresponding only to these words are updated each pass. The number and frequency in which words are selected as negative samples can be tuned (e.g., down-weighting high-frequency words).

### Training data preprocessing and word2vec training

We obtained 1,262,986 full length 16S rRNA amplicon sequences from the GeneGenes database [55]. Full length sequences were used to ensure that our embeddings were region agnostic, in that the embedding would be trained on context windows found in any variable or conserved region throughout the 16S rRNA gene, permitting query sequences to be embedded irrespective to the region of the gene they spanned. For each sequence of length N, we generated all possible subsequences (*N — k* + 1) of length *k* (*k*-mers), In natural language processing terms, these *k*-mers were treated as words belonging to a corpus, and the set of all unique *k*-mers comprise the corpus’s vocabulary. *k*-mers with degenerate bases (bases other than ACGT) were removed. Two training sets were created for *k*-mer lengths of 6 (6-mers) and 10 (10-mers).

*k*-mer embeddings were trained using gensim’s Skip-Gram word2vec implementation over 5 epochs [56]. *k*-mers occurring fewer than 100 times were removed. We varied the parameters to generate 48 model parameterizations. Using 6-mers and 10-mers, we varied the dimensionality of the embedding (64, 128, 256); the threshold in which high frequency *k*-mers were down-sampled (0.0001, 0.000001); the number of negative samples (10, 20); and the width of the context window (20, 50). Other parameters were set to their default values.

### KEGG data acquisition

16,399 full length 16S rRNA amplicon sequences were obtained from the KEGG REST server (9/2017).

### American Gut data acquisition and preprocessing

188,484,747 16S rRNA amplicon reads from the American Gut project (ERP012803, 02/21/2017) underwent quality trimming and filtering. Sequences were trimmed at positions 10 and 135 based on visualizing the quality score of sampled sequences as a function of base position [12]. Then, sequences were truncated at positions with quality scores less than or equal to 2. Truncated sequences with total expected errors greater than 2 were removed. Some American Gut samples were contaminated by bacterial blooming during shipment. Contaminated sequences were removed using the protocol provided in the American Gut documentation (02-filter_sequences_for_blooms.md). Only sequences from fecal, hand, head, and tongue body sites were kept. Head and hand were merged into a “skin” category. Any remaining samples with fewer than 10,000 total reads were removed.

### American gut OTU picking

Closed-reference OTU picking was performed with QIIME using SortMeRNA against GreenGenes v13.5 at 97% sequence identity [11]. Library size was normalized via clr, where the normalized vector of abundances 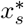 for sample s was obtained by 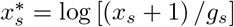, where *g_s_* is the geometric mean for sample s,

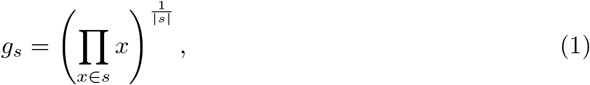

and x is the unnormalized abundance of a single OTU.

### Embedding sequences, samples, body sites, and clusters

To generate a sequence embedding, the weighted embeddings of all *k*-mers m belonging to sequence r were summed and then normalized by the total number of *k*-mers M_r_ in sequence r. Each *k*-mer was weighted based on its frequency within the query set of sequences (sequences to be embedded, but not the sequences initially used for training). Note this down-weighting is distinct from the down-weighting used during training, which down-weighted *k*-mers based on their frequency in the training (GreenGenes) sequences. Thus, we have

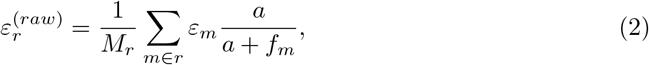

where *ε_m_* is the d-dimensional embedding for *k*-mer m, *f_m_* is the frequency of *k*-mer m across the entire set of query sequences to be embedded, a is the parameter to control the degree in which *k*-mer m is down-weighted, and *M_r_* is the total number of *k*-mers embedded into sequence r (*i.e.,* the total number of k-mers belonging to sequence r that were also present in the training set and thus have embeddings). The resulting raw sequence embedding was then denoised by removing its projection to its first principle component,

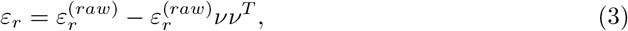

where *ν* is the first principle component obtained via singular value decomposition.

For sample, cluster, or body site embeddings, the process was instead applied to all *k*-mers belonging to all sequences from a specific sample, all sequences from a specific cluster, or all sequences from all samples from a specific body site, respectively. Note that k-mers with degenerate bases were removed (bases other than ACGT); thus, some sequences received no embedding due to no *k*-mers intersecting with the training *k*-mer embeddings.

### Cosine similarity between embeddings

For an embedding A, its cosine similarity with respect to embedding B is defined as

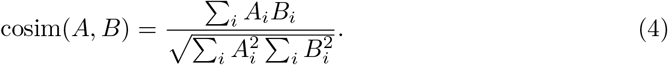

### Lower dimensional projections of the embedding spaces

For visual exploratory analysis of the embedding space, American Gut sample embeddings were reduced to 2 dimensions via t-SNE. 21 samples (7 samples each from fecal, tongue, and skin) were randomly chosen to lessen the computational burden. Principal component analysis was not performed beforehand to further reduce the dimensionality of the embedding space, as typically done. This is because the embedding space is already a lower-dimensional representation of the original input space. Centering and scaling was also not performed. Perplexity was set to 50 and t-SNE was run for 1000 iterations. sequence embeddings of the 14,520 KEGG 16S sequences were explored in the same manner. For the American gut data, because the number of total sequences was large (11,838,849), and for 10-mer embeddings in general, t-SNE was impractical in terms of time and memory requirements. Thus, to project American Gut sequence and GreenGenes 10-mer embeddings to 2-dimensions, we performed independent component analysis.

### Consensus sequence analysis

The KEGG sequences were clustered using v-search. Sequences with pairwise identity (as defined above) with its centroid below 0.8 were omitted from their respective cluster. We embedded the consensus sequence of each cluster (a consensus embedding), as well as all sequences belonging to that cluster (a cluster embedding). Then, the pairwise cosine similarities between all consensus and cluster embeddings were calculated.

### Generation of pseudo-OTUs

After generating sequence embeddings of the American Gut data, we randomly sampled approximately 1,000,000 sequence embeddings across all body sites (tongue, skin, gut) and used *K*-means [34] to cluster them into 1000 clusters (1000 pseudo-OTUs). We then obtained the centroids of these clusters. sequences from each sample were classified into the closest centroid/cluster. Finally, we quantified the number of sequences that were classified into each cluster and the abundance of each pseudo-OTU.

### Classification analysis

American Gut samples with body site (fecal, skin, tongue) labels were split into 90/10 training/testing sets containing 7526 and 835 samples, respectively. The training set was composed of 6729, 282, and 497 fecal, skin, and tongue samples, whereas the testing set consisted of 749, 31, and 54 samples, respectively. We performed multinomial classification using the lasso classifier with sample embeddings, clr-transformed OTUs, their top-256 principal components, or clr-transformed pseudo-OTUs as features. For training, we performed 10-fold cross validation to select the optimal value of the regularization parameter λ. We evaluated performance using the held-out testing set in terms of balanced accuracy, which adjusts for class imbalance by averaging the three accuracies for each individual body site:

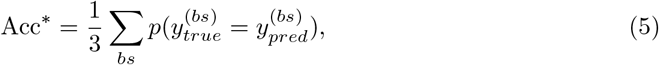

where *y_true_* is the true label and *y_pred_* is the predicated label for only samples from body site *bs.*

### Taxonomic assignment

Taxonomic assignment for both the KEGG and American Gut 16S rRNA amplicon sequences was conducted using the RDP naive Bayes classifier [57] implemented in QIIME.

### Identification of important *k*-mers

We obtained the sparse set of lasso regression weights estimated when we performed body site classification using sample embedding (described above). Each body site (skin, fecal, tongue) had its own vector of regression coefficients. To obtain *k*-mer activations, we calculated the outer product between all *k*-mer activations and the regression coefficients: *α_k_* = *ε_k_* ⊗ *β*, where, for a corpus of M *k*-mers, *α_k_* is an *M* × *J* matrix of *k*-mer activations for J body sites, *ε_k_* is an *M* × *d* matrix of *k*-mer embeddings, and *β* is a *d* × *J* matrix of regression coefficients. Each column in *α_k_* was ranked, and the top-1000 (arbitrary) *k*-mer activations for each body site were selected. For each set of 1000 *k*-mers, we identified which corresponding American Gut reads (from samples of a particular body site) contained these *k*-mers and randomly sampled 25,000 of *k*-mer-containing sequences from each body site (to ease the computational burden in the subsequent alignment step). We performed multiple alignment [58] for each set of 25,000 sequences with relatively strict gap penalties to prevent exceedingly large alignments (−25 and −10 gap opening and extension penalties, respectively). We finally mapped the position of the high-ranking *k*-mers to the alignments. The position is the range in which the *k*-mer spans in the multiple alignment, including the presence of gaps.

### Identification of important node activations

We calculated sequence activations for each body site in a similar manner as described above. To obtain sequence activations, we calculated the outer product between all sequence embeddings and the sparse set of lasso regression coefficients: *α_r_* = *ε_r_* ⊗ *β*, where, for a corpus of *R* sequences, *α_r_* is an *R* × *J* matrix of sequence activations for J body sites, *ε_r_* is an *R* × *d* matrix of sequence embeddings, and *β* is a *d* × *J* matrix of regression coefficients. Then, for a given sample, we summed each sequence activation *α_r_* to obtain a cumulative sum for all sequence activations through sequence activation *α_r_*.

Author contributions
SW selected and preprocessed the datasets, developed the embedding pipeline, trained the models, and performed the statistical analyses. ZZ performed all clustering and doc2vec analyses. JC assessed the distribution of pairwise cosine similarity. SW, ZZ, JC, GR assisted in interpretation of the results. All authors contributed to the writing of the manuscript. All authors sequence and approved the final manuscript.

